# Bayesian inference and simulation approaches improve the assessment of Elo-scores in the analysis of social behaviour

**DOI:** 10.1101/160671

**Authors:** Adeelia S. Goffe, Julia Fischer, Holger Sennhenn-Reulen

## Abstract

1. The construction of rank hierarchies based on agonistic interactions between two individuals (“dyads”) is an important component in the characterization of the social structure of groups. To this end, winner-loser matrices are typically created, which collapse the outcome of dyadic interactions over time, resulting in the loss of all information contained in the temporal domain. Methods that track changes in the outcome of dyadic interactions (such as “Elo-scores”) are receiving increasing interest. Critically, individual scores are not just based on the succession of wins and losses, but depend on the values of starting scores and an update (“tax”) coefficient. Recent studies improved existing methods by introducing a point estimation of these auxiliary parameters on the basis of a maximum likelihood (ML) approach. For a sound assessment of the rank hierarchies generated this way, we argue that measures of uncertainty of the estimates, as well as a quantification of the robustness of the methods, are also needed.
2. We introduce a Bayesian inference (BI) approach using ‘‘partial pooling”, which rests on the assumption that all starting scores are samples from the same distribution. We compare the outcome of the ML approach to that of the BI approach using real-world data. In addition, we simulate different scenarios to explore in which way the Elo-score responds to social events (such as rank changes), and low numbers of observations.
3. Estimates of the starting scores based on ‘partial pooling” are more robust than those based on ML, also in scenarios where some individuals have only few observations. Our simulations show that assumed rank differences may fall well within the ‘uncertain” range, and that low sampling density, unbalanced designs, and coalitionary leaps involving several individuals within the hierarchy may yield unreliable results.
4. Our results support the view that Elo rating can be a powerful tool in the analysis of social behaviour, when the data meet certain criteria. Assessing the uncertainty greatly aids in the interpretation of results. We advocate running simulation approaches to test how well Elo scores reflect the (simulated) true structure and how sensitive the score is to true changes in the hierarchy.

## 1 Introduction

Dominance hierarchies based on agonistic (or approach and retreat) interactions are often used to characterize the social structure of groups (e.g. Asiatic wild asses (*Equus hemionus*) (Ganslosser and Dellert, 1997), dogs (*Canis familiaris*) (Cafazzo et al., 2010), geladas (*Theropithecus gelada*) (Johnson et al., 2014), Guinea baboons (*Papio papio*) (Kalbitzer et al., 2015), Przewalski horses (*Equus ferus przewalskii*) (Tilson et al., 1988), spotted hyaenas (*Crocuta crocuta))* (Tilson and Hamilton, 1984), plains zebras (*Equus quagga*) (Ganslosser and Dellert, 1997), and sea lions (*Zalophus californianus*) (Schusterman and Dawson, 1968)). In addition, characteristics of social hierarchies such as the extent of the linearity in dominance interactions have been used in inter-specific comparisons (e.g. macaques (*Macaca* spp.) (Adams et al., 2015), and equids (*Equus* spp.) (Ganslosser and Dellert, 1997; Proops et al., 2012)).

Traditionally, scholars in the field of Animal Behaviour have generated social hierarchies by pooling sequential data into matrices, from which a quasi-static rank order can be derived for the time period under consideration (e.g. David (1987, 1988); de Vries (1998)). Implicitly, such an approach assumes that rank relationships are invariable. Due to migration events, death, or coalitionary upheaval, this assumption is rarely, if ever met (e.g. Cheney et al. (2004); Haunhorst et al. (2017)). To acknowledge changes in the rank hierarchy, one common approach was to compare the hierarchies over different periods of time (Arseneau-Robar et al., 2017), or before and after specific events, such as the immigration of a new subject (Zhu et al., 2016). Yet, this method fails to track the potentially continuous changes in the rank order of subjects. Breaking down the assessment of rank hierarchies into ever shorter windows of time is no option, however, because short sampling periods result in a lack of data to infer the rank hierarchy.

In recent years, researchers in animal behaviour studies have therefore turned to the Elo-rating method, which allows tracking the outcome of individual dyadic interactions and assessing their influence on the overall rank hierarchy (Elo, 1961, 1978; Albers and de Vries, 2001; Neumann et al., 2011). Using this method each individual is assigned a score and scores are continuously updated at each successive interaction. Comparing scores from a given time point allows the assessment of individuals’ rank relative strengths and positions. Elo-rating was initially developed to assess the rating of chess players (Elo, 1961, 1978) and has been further applied to a variety of other sports. Additional information regarding how Elo scores are generated may be found in the relevant literature: Elo (1961, 1978); Albers and de Vries (2001).

One fundamental problem with a dynamic approach, such as Elo-rating, is that the sampling begins at some arbitrary point in time where the animals in stable groups already possess (unknown) rank relationships. As a solution for studies of already existing groups, an arbitrary score is assigned to each subject at the beginning of the study (Neumann et al., 2011), akin to Elo-rating in sports (Elo, 1978). Because this arbitrary score most likely deviates from the score that would be obtained if information about the past had been available, it may lead to considerable bias in the estimates of the Elo scores, especially in the first period of the study. The period until a certain equilibrium is reached - which is itself difficult to judge in this dynamic approach - is also known as the “burn-in” phase, which is often discarded for varying reasons (Neumann et al., 2011; Franz et al., 2015).

To avoid such loss of information, Foerster et al. (2016a) recently introduced a maximum likelihood (ML) approach to estimate the starting scores, as well as the winning/losing tax constant (*k*), on the basis of the observed interactions (*x_j_*), such that the complete course of Elo scores most plausibly matches this sequence of observed interaction outcomes. This is an important contribution, since it overcomes a shortcoming of the classical way of calculating Elo scores that was depending on artificially determined starting scores, as well as an artificially determined tax constant. The ML approach is able to by-pass the problems arising from predetermined and artificial starting values (see Neumann et al. (2011)). Yet this method creates a novel ‘downstream’ problem. Specifically, in order to see whether a group of individuals already has a clearly developed hierarchy at the beginning of a study, separate models - with and without estimating starting scores - are fitted. This generates a decision task between those two models. Because decisions are always accompanied by uncertainties, these need to be incorporated in inferences drawn from the data.

Further, little is know about the elasticity of the Elo rating method. One of the benefits of this method is the possibility to assess the temporal dynamics in winner-loser interactions, yet we know very little regarding how fluctuations in Elo scores emerge and how well they reflect “true” scores. Three aspects of the temporal responsiveness of the Elo rating method are of specific importance: (1) the response to changes in the true hierarchy, (2) the response to variations in the predictability of each single outcome, and (3) the response to varying interaction rates within the group.

In order to address these concerns we suggest using an approach that: (i) gives a quantification of the variation in the starting hierarchy, (ii) shows how this interplays with the winning/losing update coefficient (*k*), and (iii) directly introduces the above “model decision uncertainty” into a single estimation result. We achieve this by a Bayesian inference (BI) approach applying the concept of *partial pooling*, which assumes that all starting scores are samples from the same distribution with a shared variation parameter σ. Partial pooling is the current state-of-the-art concept for tasks where a compromise is searched between modeling the full individual variation by independent individual-specific coefficients (no pooling, equal to the ML result), and modeling no individual variation (full pooling, only one single population coefficient) within a population: each individual is assumed to have a different ability for winning an encounter. However, the data for all of the observed individuals - collectively forming a population - inform the estimate of each individual, leading to an optimal and data-driven redistribution of estimation precision and imprecision among the individual-specific coefficients (Gelman and Hill, 2007, Chapter 12).

There are few large data sets containing long-term data which offer the opportunity to assess temporal variation in social dominance hierarchies. Here, we use real-world data from Foerster et al. (2016a) in order to compare the utility of ML estimation to our newly introduced BI approach (using partial pooling) in the prediction of starting values. This will show the the extent of uncertainty associated individuals’ with Elo scores in a large natural system. In order to delve into some of the variables which may be responsible for variations in uncertainty, we use simulations to explore issues associated with how well Elo scores 1) represent the true rank order and rank distances in unbalanced designs in which dyads vary in their interaction rates, and 2) respond to changes in the true hierarchy.

The remainder of this manuscript is organized as follows: We begin by introducing a BI approach for estimating starting scores and the Elo score winning/losing tax constant (*k*). Subsequently, we present the results of a re-analysis of real-world data, followed by simulation studies to evaluate several aspects that are important when working with Elo scores on behavioural data. The Stan (Carpenter et al., 2017b) code implementing the BI approach is included in supplementary material S1, and the full R code implemented in the statistical software environment R (RC Team, 2017) used for the purpose of this article (real-world data analysis and simulation-scenarios) is available under https://github.com/holgersr/Bayesian-inference-and-simulation-of-Elo-scores-in-analysis-of-social-behaviour.

## 2 Materials and Methods

### 2.1 Calculating Elo scores

Immediately after each interaction *j* = 1,…, *J*, the Elo scores (Elo_*i,j*_ of individuals *i* = 1,…, *n*) are recursively derived (following Franz et al. (2015)) as a function of Elo scores before the first interaction (Elo_*i*,0_), all interaction outcomes in between, and a winning/losing tax coefficient (*k*), by:

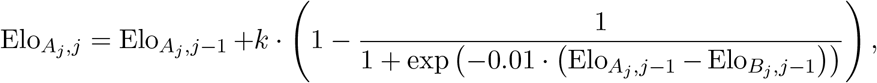

for individual *A_j_* winning over individual *B_j_* (*A_j_, B_j_* ∈ {1,…, *n*}) - the Elo score of *A_j_* increases –, and by:

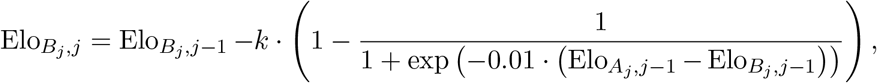

for *B_j_* losing against *A_j_*, in which case the Elo score of *B_j_* decreases. Here, *A_j_* and *B_j_* denote the two individuals participating in interaction *j*. For all remaining individuals *i ∉ {A_j_, B_j_*}, the Elo score remains unchanged, ie. Elo_*i,j*_ = Elo_*i,j*-1_. Note that for notational convenience, we will use only *A* and *B* in the following to refer to *A_j_* and *B_j_*.

In an Elo update as described by the above equations, the central role is played by the factor:

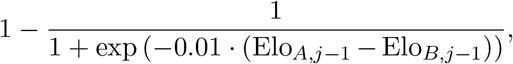

which describes the probability that *A* loses interaction *j* against *B*. This is a simpler way of writing down the logistic function, known as response function from the logistic regression model (e.g. in Fahrmeir et al. (2013), Section 5.1.1):

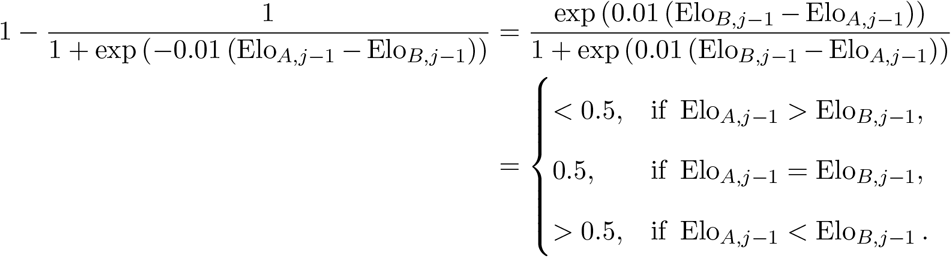

See Supplement S2 for a derivation of this equation.

By the recursive derivation of Elo scores, the Elo scores before the first interaction Elo_*i*,0_ (denoted as *starting scores* in the following) become ‘active’ at the interaction where individual *i* is interacting for the first time. So, the index terminology with 0 for the interaction index does not strictly refer to the interaction index before the first observed interaction of the whole group (i.e., it does not necessarily mean *j* = 0), but refers to the interaction previous to the first observed interaction of individual *i*.

In the approach introduced by Foerster et al. (2016a), this estimation is achieved by maximizing the log-likelihood function *l*, which is defined by the sum of log (*p_A,B,j_*) across all observed interactions *j*, i.e.

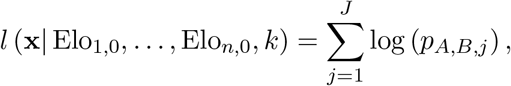

where *A* and *B* denote the individuals involved in interaction *j*, x denotes an observation vector containing all interaction outcomes *x_A,B,j_, j* = 1,…,*J*, and *p_A,B,j_* is the probability that *A* wins against *B* in interaction *j*, i.e.

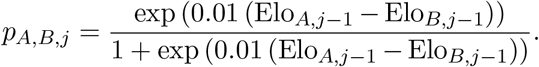

### 2.2 Reformulation of the winning/losing probability term

In the above definition of Elo scores, the equation:

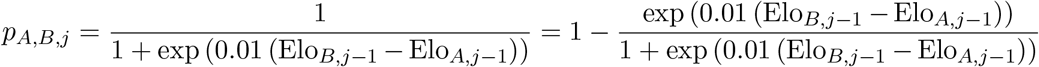

determines the probability that *A* wins interaction *j* against *B* (this is according to the definition by Albers and de Vries (2001); see also Franz et al. (2015); Neumann et al. (2011)).

We use a slightly different formulation for this winning probability equation that replaces the factor *δ* = 0.01 in the denominator by *δ* = 1:

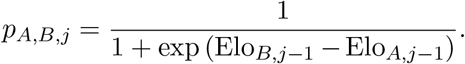

This definition is equivalent to the well-known logistic distribution function for a random variable defined by dyadic difference in Elo scores, i.e. by Elo_*B,j*-1_ – E1o_*A,j*-1_, and has some benefits for interpretation: For Elo*_A_* - Elo*_B_* measuring the difference in Elo scores between two individuals *A* and *B*, the neutral odds 1 = 1,1 for *A* winning over *B* are multiplied by exp(Elo*_A_* – Elo_*B*_), i.e.:

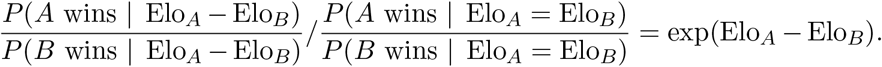

By using *δ* = 1, differences between Elo scores become directly interpretable on the familiar logistic scale: a difference in Elo scores of the value 5 - such as one individual has an Elo score of 1 and the other individual has an Elo score of 6 - leads to a probability of 99.3% (*R* command `plogis(5)`) that the individual with the higher Elo score wins an interaction between these two individuals and a probability of 0.7% to lose it. For the difference values of 0, 1, 2, …, 5, the call of the `plogis()` function returns: 50.0%, 73.1%, 88.1%, 95.3%, 98.2%, and 99.3%.

It only needs basic calculation rules for random variables to demonstrate (Supplement S3) the standard deviation equivalence by varying *δ*, and also how *k* scales with *δ*.

### 2.3 Bayesian estimation of starting scores and tax coefficient (*k*)

In contrast to Foerster et al. (2016a), we use a fully Bayesian statistical inference approach to estimate starting scores and *k* (Supplement S1 gives `Stan` code for a software implementation). With this, we are able to make probabilistic prior statements - for all of the coefficients (Elo_1,0_,…, Elo_n,0_,*k*) included in the probabilistic formulation of the data generating mechanism, i.e. included in the likelihood - in the form of probability densities:

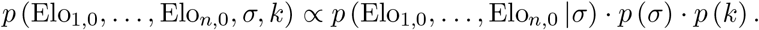

With the use of Bayes’ theorem, we may base post-estimation calculations as well as inferences for the starting scores (Elo_*i*,0_) and the tax coefficient (*k*) on a direct probability density function termed the *posterior density*:

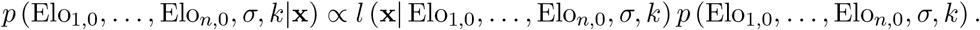

#### 2.3.1 Prior statement for starting scores

In application, individuals are observed with a different number of interactions, where it is rather the rule than the exception that some individuals only have a very small number of observed interactions. For those rarely observed individuals, only scant information is available, leading to increased standard errors in a direct ML estimation. Importantly, in extreme cases where an individual either losses or wins all interactions, the marginal likelihood is completely flat, and the starting score estimation becomes non-identifiable.

Instead of estimating the starting scores completely independently (as performed in ML estimation), a solution to the above problem is to estimate a scale parameter for a shared prior distribution of starting scores. This approach is the previously mentioned *partial pooling*, suitable in various application scenarios (Gelman et al., 2016) and conceptually very close to Bayesian Ridge Regression (Fahrmeir et al., 2010).

We choose the starting values Elo_*i*,0_ coming from a shared population distribution:

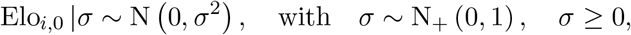

where, without any loss of generality, we fix the mean of the prior for the starting scores to 0, since the mean Elo score is not well-defined (recall that in the definition of Elo scores, the value of a score only matters relative to the other scores). For the scale parameter σ, the distribution N_+_ (0,1) denotes a truncated (beyond 0) standard normal distribution (The remainder of this passage contains practical guidelines of how to sort in and set up this prior adequately).

This “shared-distribution-assumption” acts as an informative prior to the starting scores, where a general rule of Bayesian statistics applies that the strength of a prior is relative to the flatness of the marginal likelihood: for the influence of a prior, it only matters how flat the prior is in the region of high likelihood, and vice versa.

- For an individual with a compact region of high marginal likelihood (an ‘‘informed individual”), the prior has a very small gradient across this range, the prior has only a weak influence.
- For an individual with a wide region of high marginal likelihood (an “un-informed individual”), the prior incorporates its full shape, the prior consequently has a strong influence (since the mean of the partial pooling prior is equal to the mean of the starting scores, this prior has a regularizing influence). This mechanism has the consequence that the estimate of the individual becomes biased towards 0, but - for only barely informed individuals - we want to have this bias, as it works stronger on extreme quantiles of the coefficients distribution in comparison to it’s mean, i.e., the corresponding standard error is estimated much more realistically by the price of a (usually small) bias.

By this, partial pooling cures the problem of inflated standard errors in a very natural way (“at a cost of (typically small) bias”(Fahrmeir et al., 2013, p. 238)): when the knowledge about an individual is very sparse, the most plausible assumption is to assign it a value that does not differ substantially from the population mean, i.e. the individual is ranked in the center of the hierarchy, and does not stand out as either very dominant or subordinate. Partial pooling thus allows a single starting score to speak with a louder voice if it is based on strong information, i.e. a large group of observations from the same individual. Conversely, if little information is available, the starting score does not have a strong influence.

The use of the logistic distribution function is again helpful to interpret the scale parameter σ in the distribution of starting scores: σ is the standard variation of a normal distribution, and [-2 · σ, 2 · σ] roughly spans an inner 95% probability interval of starting scores. Applying the logistic quantile function, we can transform this to an inner 95% probability interval for winning probabilities in comparison to an individual that is expected to win against the one half of the group, and to lose against the other half; such an individual would have the central Elo score of 0. This view on σ from the perspective of assumed differences in Elo scores, and the differences in probabilities they refer to, acts as a helpful tool while thinking about priors for σ.

Although the use of a truncated standard normal distribution prior for σ is not uncommon, we want to point out that, as for any prior assumption in practice, sensitivity checks should be performed. For example by the use of Hellinger distances in the spirit of Roos and Held (2011) or by refitting using a different prior formulation (see Figure 3 for an example).

#### 2.3.2 Prior statement for winning/losing tax coefficient

The winner and loser effects commonly exist in social groups and may interact with each other in order to generate the social dominance hierarchy (Dugatkin, 1997; Neumann et al., 2011). The winning/losing tax coefficient (*k*) plays a central role in allowing for - and measuring - dynamics of hierarchies by Elo scores.

Albers and de Vries (2001), as well as Neumann et al. (2011) used different selections for *k* in order to compare the changes in Elo scores and hierarchies that go with it; traditionally, these *k* values are predetermined and fixed between 16 and 200 (e.g. Neumann et al. (2011)).

Foerster et al. (2016a) bound exp_*e*_(*k*) between -10 and 10 (in the dryad archive they refer to in their article), and by this *k* between approximately 0.000 045 and 22 000. This shows that they allow a much wider spread to *k* than Neumann et al. (2011), but certainly not all of those values make, a priori, equal sense.

This can be demonstrated using two toy-examples:

- if *k* is 1, and two individuals *A* and *B* with scores Elo*_A_* = 1 and Elo_*B*_ = 3 (*δ* = 1) have an encounter (`plogis(2)` is 0.88), then the scores change to Elo_*A*_ = 0.88 and E1o_*A*_ = 3.12 if *A* wins, and to 1.88 and 2.12 if *B* wins. The post encounter probability (that the originally higher scored individual *B* wins) is 0.90 in the first case, and 0.55 in the second case.
- everything unchanged, but now *k* = 2 instead of *k* = 1, the scores change to 0.76 and 3.24 if *B* wins, and to 2.76 and 1.24 if *A* wins. The post encounter probability (that the originally higher scored individual *B* wins) is 0.92 (`plogis(3.24-0.76)`) in the first case, and 0.18 (`plogis(1.24-2.76)`) in the second case.

Thus, the selection of *k* = 2 (for *δ* = 1) results in a complete turn-around of the two scores in the case of the unexpected outcome. Consequently, any larger values will hardly yield sensible results.

Because of this, we attribute the same prior to *k* as we have done to σ:

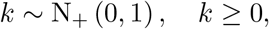

where N_+_ (0,1) again denotes a truncated (beyond 0) standard normal distribution. This informative prior is ideally strong enough to avoid pathologically large posterior samples for *k*, and still weak enough to “let the data speak for themselves”. Again, sensitivity checks to the prior choice are recommended in practice.

#### 2.3.3 The effect of newly introduced members on the ranking of ‘‘silent members”

In the analysis of long-term data on agonistic interactions of female eastern chimpanzees (*Pan troglodytes schweinfurthii*) (Foerster et al., 2016a), individuals exiting (through emigration or mortality) typically had achieved high Elo scores, while newly entering individuals (through immigration or birth) had low Elo scores. Remaining individuals who had not been involved in any interactions for a long period of time (we call them “silent members”), automatically “march through” the hierarchy (see Figures 1 (e, f) and 2 of the female eastern chimpanzee data from Foerster et al. (2016a)).

**Figure 1:**
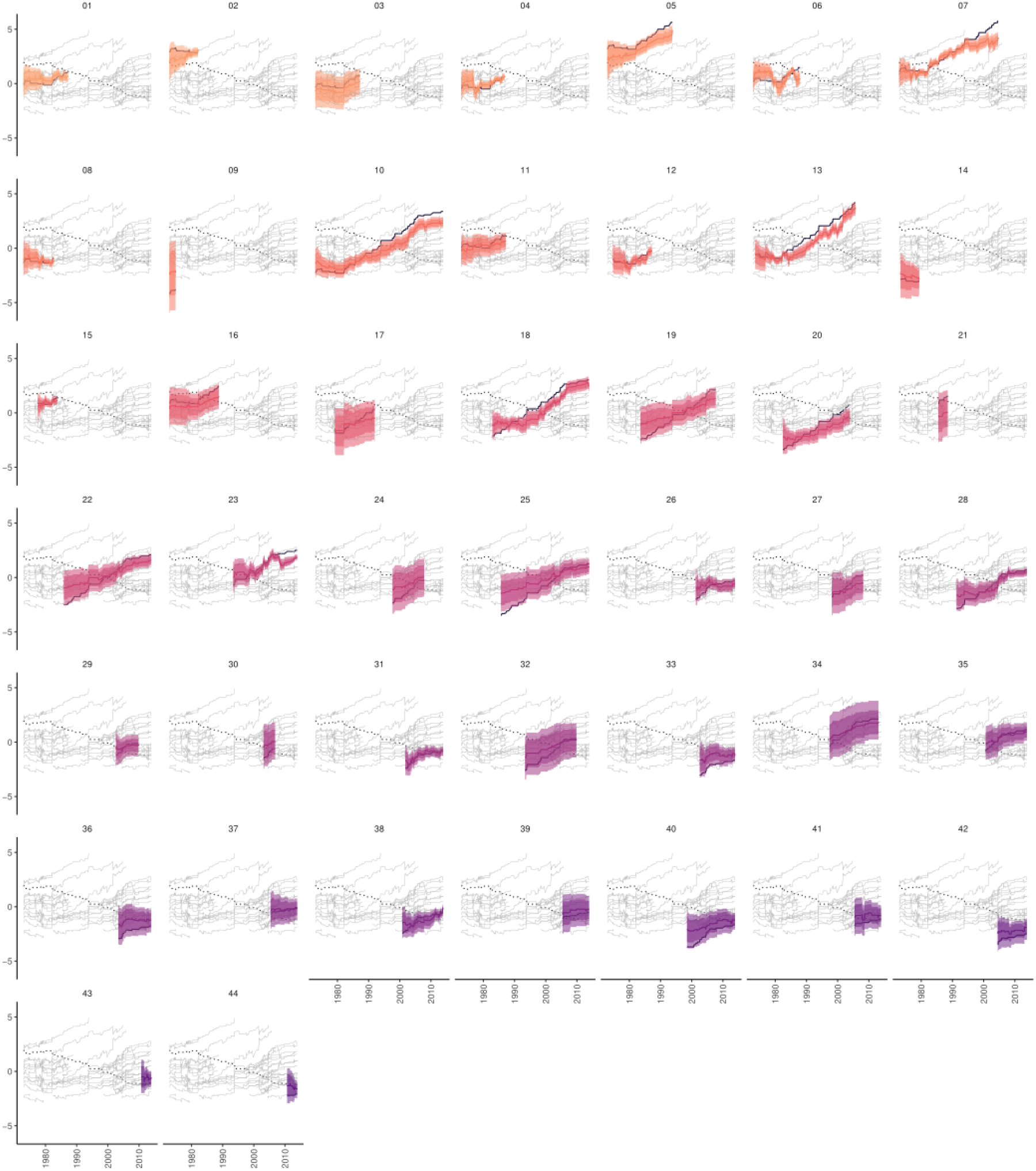
Estimated Elo scores for the Foerster female data (x-axis: time in years; y-axis: Elo scores): Posterior means of our Bayesian estimation approach are shown as solid coloured lines (step functions), results from Foerster et al. (2016a) as solid black step functions. Shaded areas show 95% and 80% credible intervals, incorporating the uncertainties about starting scores as well as about *k*.

**Figure 2:**
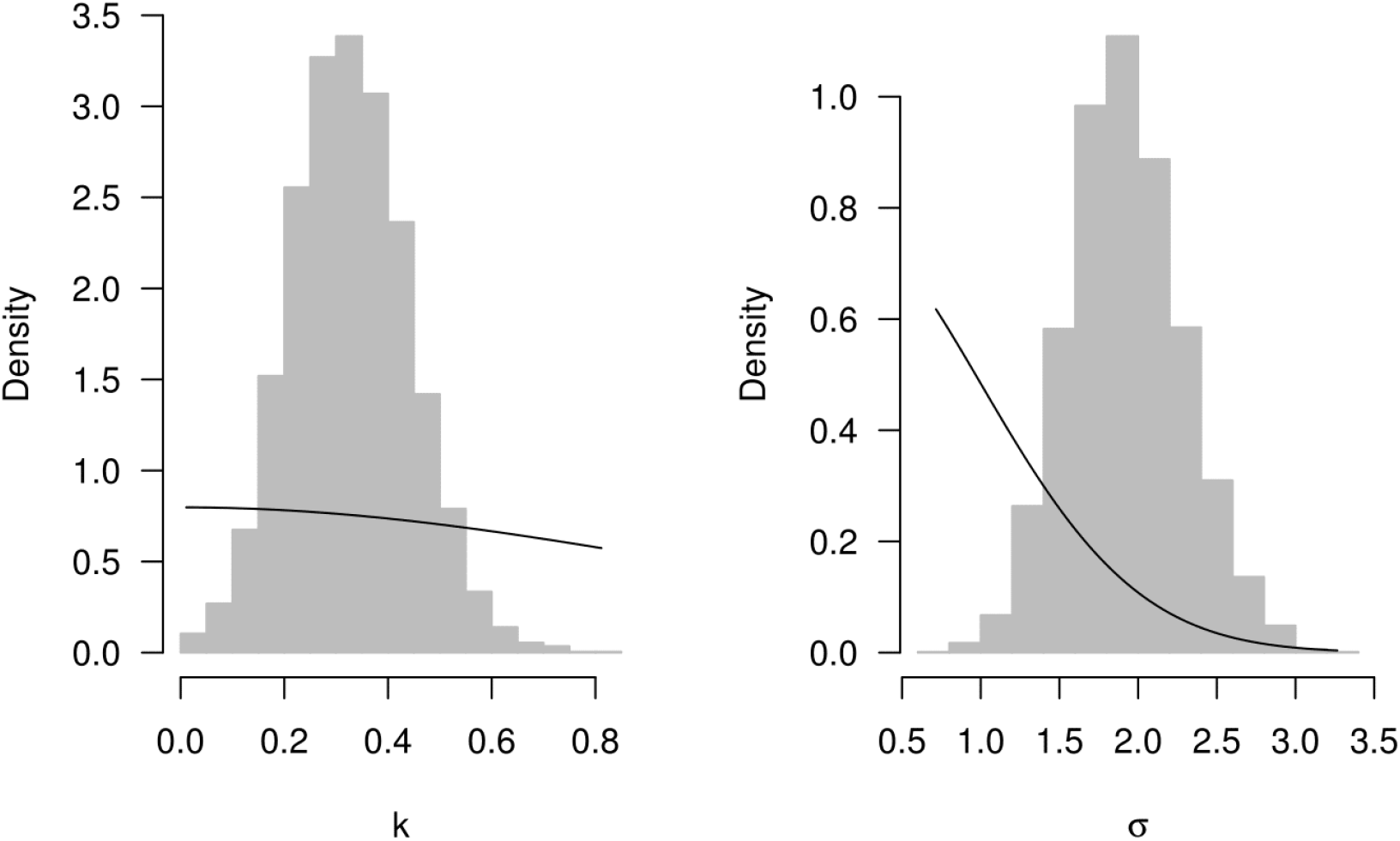
Comparison of posterior density (grey shaded histograms) to the prior density (solid black line) for tax coefficient *k* ~ N_+_ (0,1), and starting score standard deviation σ ~ N_+_ (0,1) (Note the small scale of *k* because of the reformulated winning/loosing probability term with *δ* = 1; see Section 2.2).

We therefore decided to subtract the mean Elo score of the present individuals at each interaction from the Elo scores of the present individuals at this interaction (i.e. not-present individuals do not change), as per Franz et al. (2015). Each individual’s Elo score is then directly interpretable - at any time - as the difference to the mean of the current group, which is 0. By this, changes in Elo scores of “silent members” become more clearly visible. This ‘‘correction” of the mean is also important in the Bayesian estimation of starting scores, since the applied partial pooling approach assumes that all starting scores are sampled from a distribution with equal mean, which we need to guarantee in the algorithm (see Supplement S1).

### 2.4 Real-World Application

We first conducted a re-analysis of long-term data (Foerster et al., 2016b) on agonistic interactions of female eastern chimpanzees, studied at Gombe National Park, Tanzania. The study population consisted of *n* = 44 female individuals and data were collected between the years 1969 - 2013. As in Foerster et al., we did not include the first 100 interactions in order to facilitate comparison between the ML and BI approaches; a total of *J* = 915 agonistic interactions were included in the analysis.

This dataset was ideal as it consists of detailed information regarding social relationships over an extended time period in a long-lived, group-living species. This dataset also allowed us to study the ML and BI estimation approaches in a scenario in which one individual contributes with losses (or wins) only. The occurrence of only wins or losses is inadvertently the case when extremely strict (despotic) and completely stable hierarchies (potentially with low interaction rates) are analysed. Foerster et al. (2016a) addressed this issue by removing 13 females that incorporated solely wins or losses in order to ‘‘facilitate model fitting and interpretation” (p.2).

As a further in-sample prediction criterion for this application, we calculated mean Brier scores (Brier, 1950):

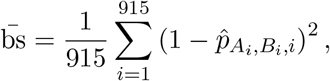

which not only incorporates the correct sign of the Elo score difference, as does the correctly predicted interaction proportion, but further takes into account whether the Elo scores were able to clearly distinguish the winning from the losing individuals by a (considerably) large difference in their winning probabilities. A large positive difference is a greater success in comparison to a small positive difference; a small negative is not as bad in comparison to a large negative difference.

For a quantification of the ability of Elo scores to correctly rank the individuals at each time point *j*, we calculated the mean absolute error (Ranks MAE) in relation to the underlying true ranks for the different simulated data scenarios:

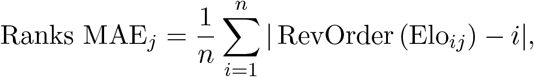

where the RevOrder (for ‘reversed order”) function gives a value of 1 to the highest Elo score at the current interaction index, and a value of *n* to the lowest Elo score (note that this definition only works if prior to calculation the individual index *i* = 1,…,*n* already gives the rank in the truly underlying hierarchy).

### 2.5 Build up for simulation 1-High-dimensional ‘‘estimation problem”

We simulated different numbers of interactions in the scenario given by the Foerster et al. (2016a) female data (*N* = 44, *k* = 0). The marginal probability that an individual is involved in one of the simulated interactions was equal for all the individuals, which should result in a more balanced data-set as used in Foerster et al. (2016a). This was achieved by randomly sampling one individual for each of the simulated interactions. The opponent identity was realized by sampling from the respective remaining individuals, with the probability being inversely proportional to the true underlying difference in starting scores to the already sampled individual (by this, dyads closer in hierarchy interact more often than dyads more distant in hierarchy). The true underlying starting scores were sampled according to our result on the female eastern chimpanzee data (Elo_0,*i*_ ~ N (0,σ^2^ = 2.62^2^), when fixing *k* to 0). Since there were no changes in the underlying scores estimated in the original study, a true tax coefficient of 0 was utilized in this simulation scenario. The numbers of interactions were 250, 500, 1000, 1500.

### 2.6 Build up for simulation 2-General unbalanced design

In comparison to the above simulation, the marginal probability that an individual is involved in one of the simulated interactions was proportional to the true underlying scores, i.e. the lower the score, the lower the interaction rate. We reduced the number of individuals to *N* = 10, to obtain a clearer picture of the uncertainties in this scenario. To introduce a tax coefficient different from 0, we swapped the scores of neighboring ranks (after 2000 interactions) as 2 with 1, 4 with 3, …, 10 with 9. We further based this simulation on an equidistant grid of underlying Elo scores from -6 to 6 with a length of *N* = 10. This simulation was repeated twice, which led to individual specific numbers of interactions between 398 and 1243 in the first run, and between 410 and 1207 in the second run.

### 2.7 Build-up for Social Instability Simulations

In order to assess the flexibility of Elo rating to adapt to changes in social dynamics, we conducted three further simulations representing scenarios similar to social hierarchy changes which may naturally occur in animal groups. All estimations were conducted using the BI approach, with assessing statistical instability by calculating the proportion of mismatches.

#### 2.7.1 Simulation 3A-External Takeover

In order to simulate a change in social hierarchy which might occur in the event of an external rank takeover, for instance as a result of an immigration event (Marty et al., 2015; Cheney et al., 2004), we used an equidistant grid of underlying Elo scores from −6 to 6 with a length of *N* = 10. The simulation was divided into two periods. In the first period of 2000 interactions we included 10 individuals and a stable social hierarchy. In the second period of 2000 interactions, the formerly dominant individual dropped to the bottom position (Elo score of 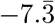), and another individual (the ‘‘immigrant”) is introduced at the top (Elo score of 6), taking over the alpha position. During this second period, the underlying social hierarchy was also stable.

#### 2.7.2 Simulation 3B-Coalitionary Leap

In the second scenario we generated a dataset which represented a coalitionary-based rank change, such as one that might occur in a nepotistic society (Weiß et al., 2011). In the first period of 1000 interactions we included 25 individuals, divided into five subgroups of five individuals each, with a stable social hierarchy. In the second period of 2000 interactions, the third ranked group was moved to the top position; for the sake of simplicity we did not introduce a change in position within the subgroups. We based this simulation on an equidistant grid of underlying Elo scores from -6 to 6 with a length of *N* = 25.

#### 2.7.3 Simulation 3C-Mortality-Instability-Recovery

In the third scenario we mimicked a mortality event followed by social upheaval and subsequent stability (Kaburu et al., 2013). In the first period of 1000 interactions we included 10 individuals in a stable - meaning no changes throughout the whole period - social hierarchy (equidistant grid of underlying Elo scores from −6 to 6). In the second period of 1000 interactions, the top two ranked individuals were removed from the hierarchy and the ranks of the remaining 8 individuals were unstable: after every 100 interactions, the underlying true Elo scores are randomly permuted. In the third period of 1000 interactions the social hierarchy was again stable.

## 3 Results

### 3.1 Real world application: female agonistic interaction data

Bayesian estimation indicated that the use of a single estimation run (consisting of similar starting values and a small/large *k*) was more appropriate than the two-step model comparison approach: posterior draws for σ ranging between 0.5 and 3.5 clearly show the uncertainty in this quantity, which is not represented in the final estimates by a selection between two point values (see Figure 2). By the use of the BI approach applying partial pooling, the increased uncertainty - introduced by selecting among different models - is directly transfered into the estimation problem.

The uncertainty of Elo scores - as illustrated by the shades in Figure 1 - indicated a considerable degree of uncertainty in the estimated coefficients. When we calculated correctly predicted interaction proportions for BI and ML (Foerster et al., 2016a), we yielded 90.7% (BI) in comparison to 89.4% (ML), respectively (we were not able to reproduce the value of 89.8% as given in Table 1 in Foerster et al. (2016a)). In comparison to the ML approach the BI approach led to a reduction in Brier score from 0.085 to 0.075.

Bayesian estimation also allowed us to achieve our primary aim of reliably estimating Elo score estimates. This came directly from the distribution/spread of posterior samples. What seemed to be a clear hierarchy for most of the individuals (Foerster et al., 2016a, Fig. 1f), appears in fact indistinguishable: there is a high degree of overlap in credible intervals and Elo-score estimates between individuals, in especially in the centre of the hierarchy (Figure 1). Solely concentrating on the Elo score point estimates, i.e. the grey lines in Figure 1, the visual comparison of the differences between the ML and the BI approach did not suggest fundamental consequences at the practical level, with only a slight advantage for the BI approach as given by the above results for Brier score and correctly predicted interaction proportions.

The BI approach did not yield strong evidence for the claim *k* = 0 (Figure 2).

### 3.2 Simulation results

#### 3.2.1 Simulation 1-Maximum likelihood vs Bayesian estimation in a high-dimensional design

As illustrated by the results in Figure 3, the estimation based on the BI approach led to an improved stability in comparison to the ML approach. Even for relatively high numbers of observed interactions, the ML estimation led to an unreliable estimation of point estimates that differed substantially from their underlying ground truth (dashed line).

**Figure 3:**
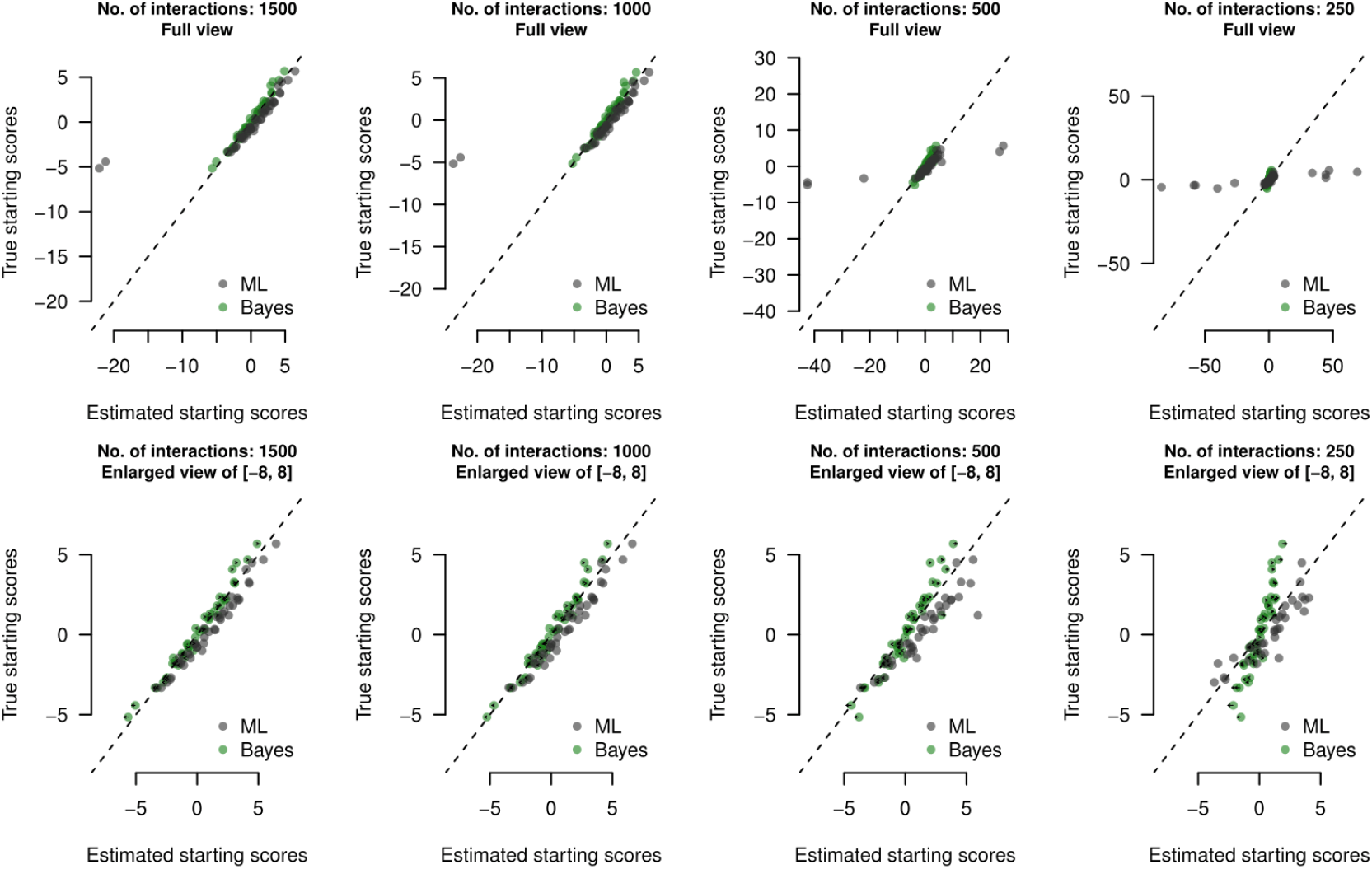
Results of starting score estimation for simulated data: y-axes values are true underlying starting scores, x-axes show estimated starting scores (posterior mean for Bayesian approach, ML estimates for the maximum-likelihood approach). Black arrows show the influence of changing the prior from σ ~ N_+_ (0,1) to σ ~ N_+_ (0, 2): For small sample sizes, a larger prior standard deviation lead to a slight reduction in the bias of the estimates, for larger sample sizes, the posterior is more and more dominated by the Likelihood, and therefore the prior influence vanishes.

#### 3.2.2 Simulation 2-General unbalanced design

If the number of interactions is highly unbalanced, it is likely that the estimation of the Elo scores of individuals with low interaction rates were inaccurate. In the present scenario, individuals with lower rank interacted at lower rates than more dominant individuals. For individuals with comparatively low sample sizes, Elo scores appeared to be more volatile and the lower portion of the dominance hierarchy was unstable (posterior means are more close to each other, credible intervals considerably overlap; Figure 4).

**Figure 4:**
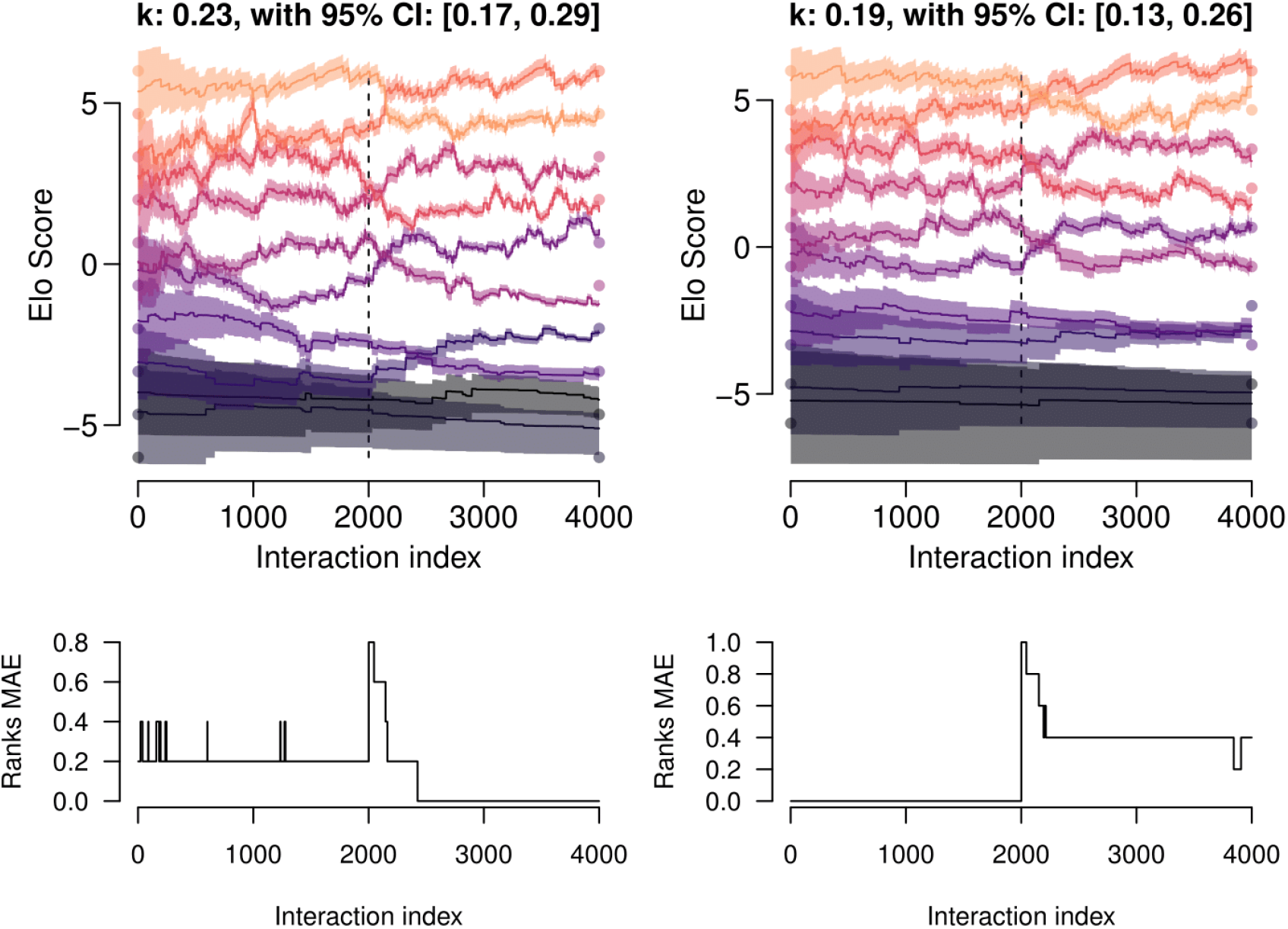
Results of two runs for simulation 2. Shaded areas give 95% posterior probability intervals for the Elo scores at each interaction, based on the posterior samples for the starting values. Filled circles at the outer extremities of the two periods indicate the individuals’ underlying Elo scores. The bottom panels show the evolution of the ranks’ mean absolute error (MAE) across the 4000 interactions.

#### 3.2.3 Response of Elo scores to changes in the hierarchy

##### Simulation 3A-External Takeover

Figure 5 shows the results of two different simulation runs in which an external takeover of the alpha position occurs: At 200 interactions after the external takeover, Elo scores began to reflect the true underlying rank order. There was consistent underlying stochasticity in both stable periods.

**Figure 5:**
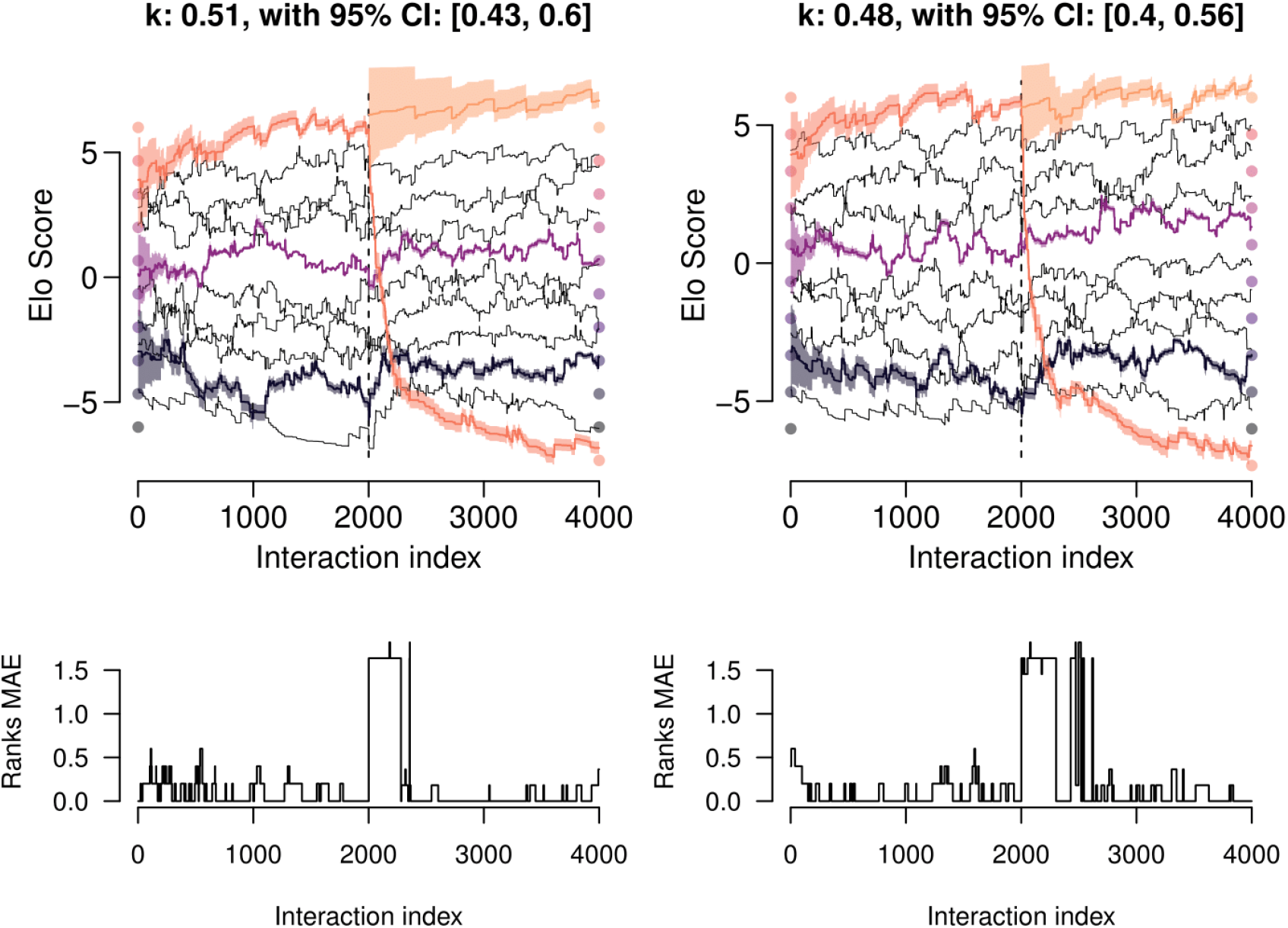
Results of two runs for simulation 3A (‘external takeover’). In the top panels the two periods are separated visually by a vertical dotted line; ten individuals are present in the first period and eleven individuals are present in the second period. The Elo scores of three individuals are highlighted by colors, and their 95% posterior probability intervals are shown. Filled circles at the outer extremities of the two periods indicate the individuals’ underlying Elo scores. The bottom panels show the evolution of the ranks’ mean absolute error (MAE) across the 4000 interactions.

The degree of uncertainty varied during different periods of the interaction sequence (Figure 5). Following the BI approach, uncertainty, as indicated by credible intervals, was initially higher in all 10 individuals at the start compared to the end of period 1. In period 2, the uncertainty of the new alpha was highest while the uncertainty of all ancillary individuals was reduced.

##### Simulation 3B-Coalitionary Leap

Figure 6 shows the results for two different simulation runs: Although order and position remained constant within the 5 subgroups, the calculated Elo scores showed a high degree of stochasticity even 2000 interactions following the change in underlying scores.

**Figure 6:**
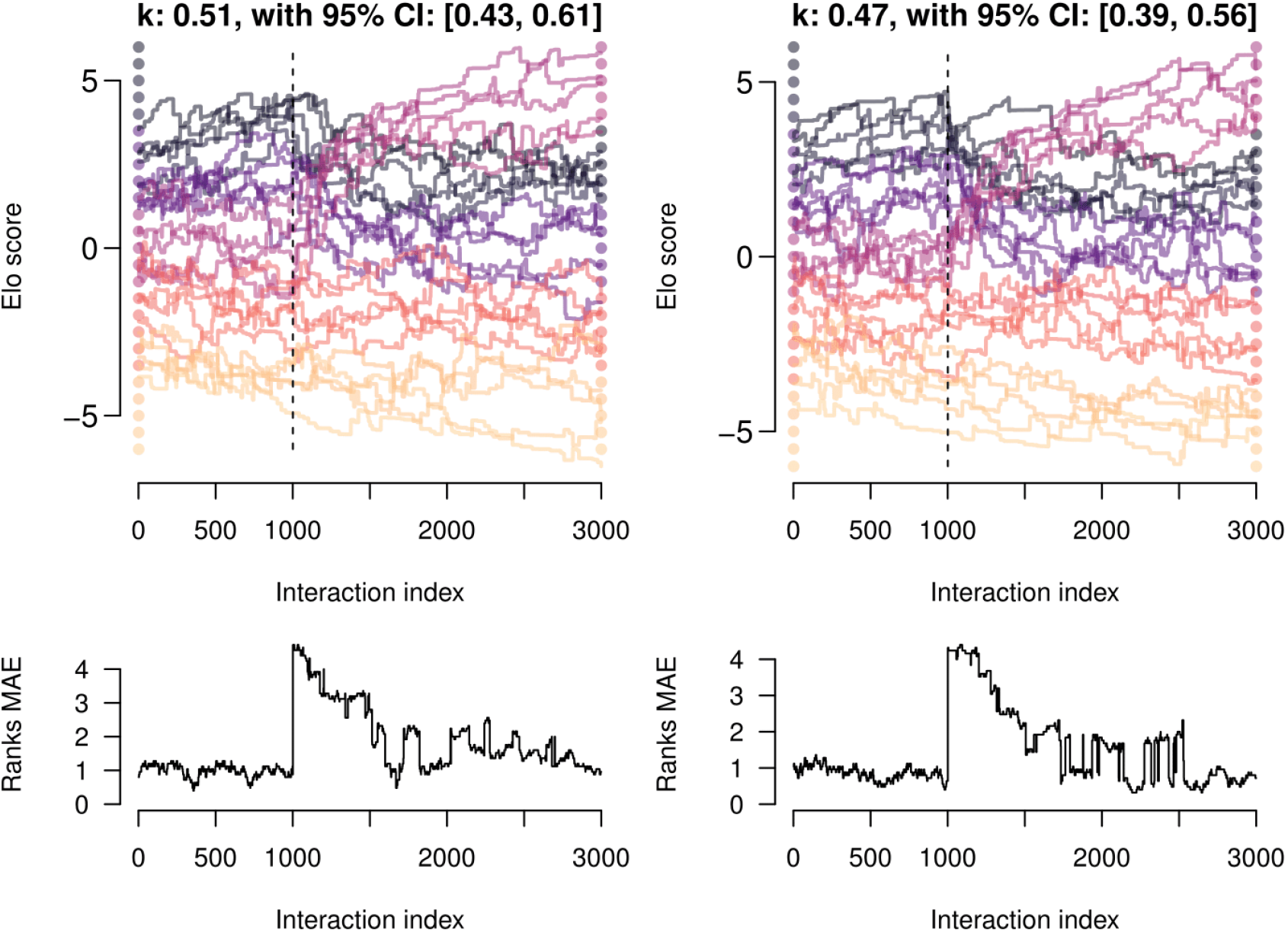
Results of two runs for simulation 3B (‘coalitionary leap’). Filled circles at the outer extremities of the two periods indicate the individuals’ underlying Elo scores and the dashed vertical line separates the two periods. The bottom panels show the evolution of the ranks’ mean absolute error (MAE) across the 3000 interactions.

##### Simulation 3C-Mortality-Instability-Recovery

Figure 7 shows the results for two different simulation runs depicting such a “mortality-instability-recovery” scenarios: in spite of a prolonged period of social disruption, Elo scores (and the mean absolute error) returned within about the first 100 interactions to the levels before the event.

**Figure 7:**
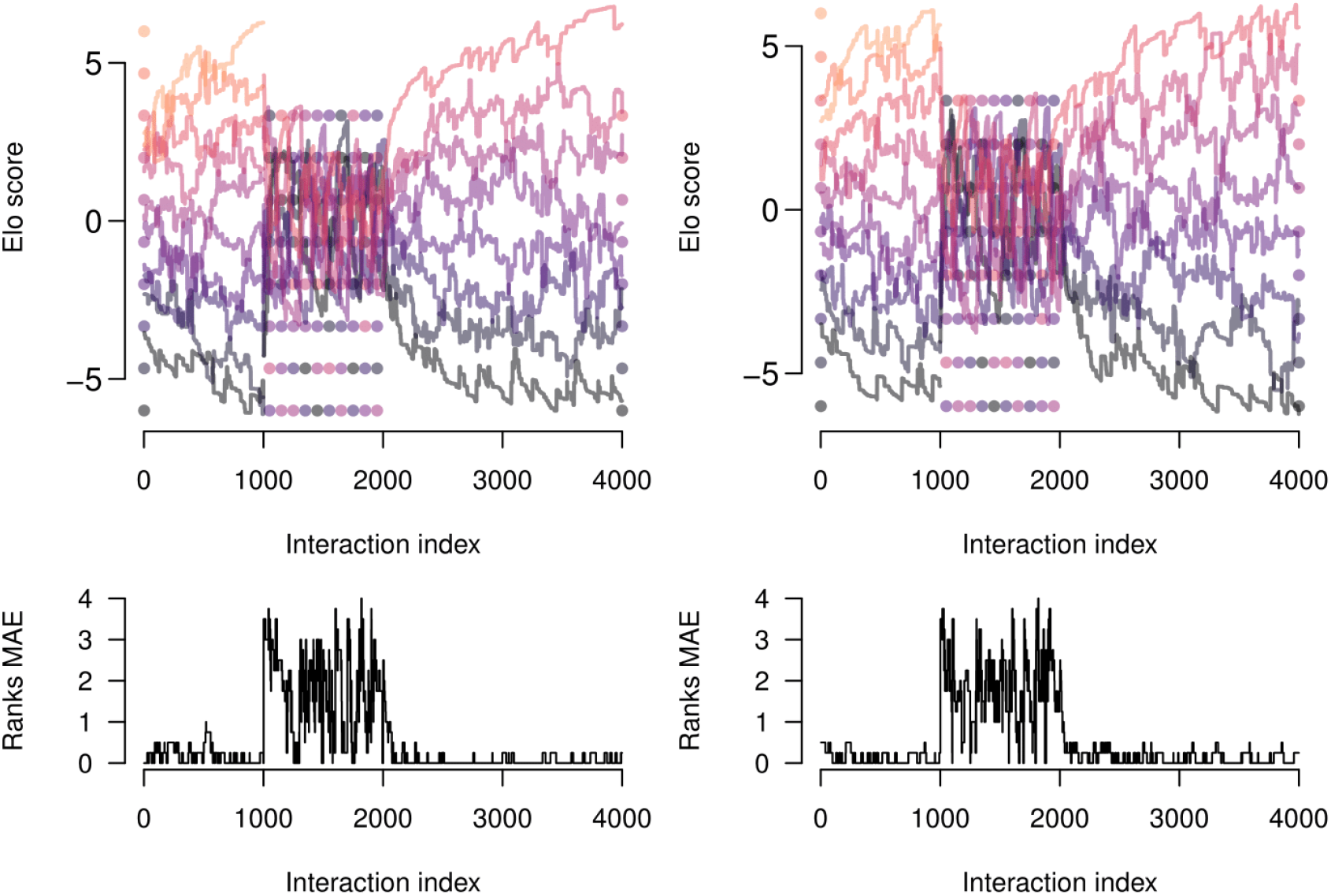
Results of two runs for simulation 3C (“mortality-instability-recovery”). Filled circles indicate the individuals’ truly underlying Elo scores. The bottom panels show the evolution of the ranks’ mean absolute error (MAE) across the 4000 interactions. The Ranks MAE calculation only incorporates the 8 individuals present in all three periods.

## 4 Discussion

Elo scores allow for the generation of dominance hierarchies while considering temporal dynamics and sequential events (Neumann et al., 2011). These scores therefore add the critical temporal dimension to the analysis of agonistic interactions between individuals. Our research extends previous applications in three critical ways: Firstly, we demonstrate that partial pooling within a Bayesian Inference approach provides better estimates of the starting scores than a ML approach. Secondly, the BI approach provides valuable additional information with regard to the assessment of uncertainty of the estimates. Third, simulation approaches allowed us to investigate the effects of different boundary conditions on Elo scores. Specifically, we found that (not surprisingly) in an unbalanced design, a small sample size increased the uncertainty of the estimate; that it may take a substantial amount of interactions (here: *N* = 200) after an external takeover until the hierarchy can be interpreted in a meaningful way; and that coalitionary leaps of whole groups may create havoc in the estimation of Elo scores, as it may take an excessive amount of interactions (in our simulation: 2000) until the hierarchy could be inferred with certainty. Finally, we found that the removal of an individual followed by a phase of instability may recover quickly.

Partial pooling appeared to be clearly superior to separate coefficient estimation. One compelling applied example is the ranking of baseball players by their estimated batting abilities (Carpenter et al., 2017a). In application to Elo scores, BI generally yields more robust results than the ML approach for the assessment of starting score point estimates - as for example shown in Figure 3 -, there was no substantial advantage of one method over the other in terms of the point estimates. Because the BI approach provides additional important additional information with regard to uncertainty, we suggest that this approach should be preferred.

The application of Bayesian estimation to a previously published data-set (Foerster et al., 2016b) allowed us to determine the extent of uncertainty in point estimates generated from real world social interactions. The assessment of uncertainty is critical when drawing conclusions about current rank hierarchies and social dynamics. For instance, what may look like a clear dominance hierarchy at a certain time point when only point estimates are considered, may provide only weak evidence for a clear underlying linear hierarchy when the estimates’ uncertainties are additionally taken into account. In such cases, it is safer to abstain from the classification of individuals with regard to specific ranks.

The simulation scenarios allowed us to assess the consequences of specific events, while having full knowledge about the underlying social interactions. In our simulations, we found various causes for uncertainty in the reliability of Elo scores. When the true values were known, Elo scores varied with the proportion/number of interactions individuals contributed to the data-set (balance), the occurrence of social change (stability), and the type of change that occurred (e.g. number and position of individuals leaving the hierarchy). As in many other cases, the number of interactions considered is critical. Importantly, even when interactions were equally balanced and hierarchies were stable, it may take large numbers of interactions in order for Elo scores to approach the “true” underlying scores. Although larger sample sizes generally seem to be preferable, we are unable to give general recommendations regarding the number of interactions in relation to the number of dyads. We do, however, recommend using a simulation approach that is based on the data available to check the sensitivity and stability of the results.

### 4.1 Inclusion/exclusion of data

In the literature, it appears that individuals with comparatively sparse numbers of interactions may be excluded when assessing dominance hierarchies (e.g. Seyfarth et al. (2012); Foerster et al. (2016a)). This strategy may potentially bias the results since the number of observed interactions per individual may be depend on the underlying hierarchy position, resulting in apparent instability in certain regions of the dominance hierarchy (for example, simulation 2).

Moreover, the exclusion of an individual removes the information from winning/losing of each of this individual’s opponents. Although ML approaches dictate the exclusion of subjects with one type of interaction solely because models could not be fitted otherwise, this is highly problematic. Effectively, if one was interested in a society with a subject that was in the top position throughout the study period (no losses), this individual would have to be excluded, and valuable information would be lost. This may even lead to a chain reaction where the second ranking subject whose loses against the former dominant subject are now removed from the analysis, also needs to be removed because it now only wins, too.

All information, seemingly minute in detail, adds to the knowledge of social dynamics in the group and can thus assist in illuminating hierarchical fluctuations. The BI approach, which is able to deal with wins/losses only situations, clearly is the method of choice under such circumstances. We therefore discourage the exclusion of subjects, or at least strongly recommend comparing results of analyses including and excluding this/these subjects, to assess the consequences of this step.

### 4.2 Uncertainty informs about the characteristics of the society

The extent to which we see uncertainty in Elo scores adds additional knowledge to our perception of the characteristics of a society. For instance, Barbary macaque (*Macaca sylvanus*) males have a rank hierarchy with clearly identifiable top and bottom subjects, while the rank for animals ranging in the middle is harder to discern (Henkel et al., 2010). For these animals, the uncertainty with which the rank can be estimated reflects precisely this fact, which would be misrepresented by a linear rank hierarchy only. Uncertainty estimates allow us to assess both the changes in rank order and the overlap of individuals with adjacent rank. In some societies, particularly those which are more egalitarian, the attribution of a fixed rank position to each individual may not be realistic. The best solution for such an egalitarian society (in which ranks are associated with such large uncertainty) might be to give up the concept of a linear hierarchy, which is a main assumption of the Elo score approach (and an egalitarian society might simply might not meet these assumptions).

### 4.3 The sensitivity of the Elo method and its drawbacks

On the one hand, Elo rating is sensitive and able to detect changes in social hierarchies resulting from social instability and demographic change. On the other hand, individual scores may be highly volatile, even in scenarios in which the hierarchy is actually stable (e.g. the first periods in Figures 5, 6, and 7). Likely, this is due to the fact that individual A’s estimated scores are indirectly affected through the interactions of individual A’s social partners (e.g., through interactions between individuals B and C). Although this allows us to pool social knowledge, and gain information about individuals with low interaction rates, the inherent stochasticity in the Elo scores of others may increase the stochasticity of third parties (in our example, subject A). Future studies should examine to which extent group size and transitivity influence the sensitivity of Elo.

With the use of different simulation scenarios, we demonstrated that it takes time for a change in the underlying hierarchy to be reflected in the Elo scores. This may depend upon the extent to which the system is already established, and the volatility of the system/degree of hierarchy stability.

### 4.4 Summary and conclusion

Elo rating has been developed in the sphere of human sporting competition in which we can watch the contests between individuals and is a tool which has recently been applied to assessing dominance hierarchies in non-human animals. However, some of the challenges we face as those who study animal social dominance are a direct result of the type and complexities of the contests we observe and the limited window by which they are viewed. Thus far, Elo rating has proven to be a powerful tool in our understanding of temporal fluctuations in dominance hierarchies. However, the role of group size, interaction rate, sample size (i.e. number of interactions) and temporal order of events remain unclear. Here, we have taken the first steps in trying to understand these factors by presenting a tool, Bayesian inference, which allows us to see the variability in uncertainty around Elo scores and some of the ways in which uncertainty can vary in simple simulated dominance hierarchies.

## Acknowledgement

We like to thank Mathias Franz and Roger Mundry for helpful comments and suggestions on a previous version of this manuscript.

## Author contributions statement

ASG and HSR conceived the ideas and designed methodology; ASG and HSR analysed the data; ASG and JF led the writing of the manuscript. All authors contributed critically to the drafts and gave final approval for publication.

## S1 Stan implementation of Bayesian estimation of starting scores and *k*

The following `Stan` (Carpenter et al., 2017b) code implements the Bayesian estimation of starting scores and winning/losing tax coefficient *k* as introduced by this manuscript. For convenience, the observations **x** were stored in the form of two vectors **A** and **B**, each of length *n*, where the integer values in **A** store the index of the interaction-specific winning individual, and the integer values in **B** store the index of the interaction-specific losing individual. The remaining input arguments (see the `data` block) are the number of encounters `N`, the number of individuals `K`, a response vector `y` always taking on value 1 (needed for the probabilistic formulation of the likelihood as a Bernoulli random variable; always 1 since it’s always the individual from `Ai` winning the encounter), and the Elo score difference factor `delta`.

~~~
functions {
 real[] ProbFunction(real[] EloStart, real k, matrix presence, int N, int K, int[] Ai, int[] Bi, real delta) {
   real result[N];
   real toAdd;
   vector[K] EloNow;
   for (j in 1:K) {
    EloNow[j] = EloStart[j];
   }
~~~

~~~
   for (i in 1:N) {
      // centering:
      EloNow = EloNow - dot_product(row(presence,i),EloNow)/sum(row(presence,i));
      // likelihood contribution:
      result[i] = 1/(1 + exp(delta * (EloNow[Bi[i]] - EloNow[Ai[i]])));
      // update addend:
      toAdd = (1 - result[i]) * k;
      // update:
      EloNow[Ai[i]] = EloNow[Ai[i]] + toAdd;
      EloNow[Bi[i]] = EloNow[Bi[i]] - toAdd;
      }
   return result;
 }
}
~~~

~~~
data {
   int<lower=1> N; // number of encounters
   int<lower=1> K; // number of individuals
   int<lower=1> Ai[N]; // winner’s index
   int<lower=1> Bi[N]; // losers’s index
   matrix[N, K] presence;
   int<lower=0> y[N]; // always 1
   real<lower=0> delta; // Elo Score difference factor
}
~~~

~~~
parameters {
   real EloStart_raw[K];
   real<lower=0.0> k_raw;
   real<lower=0.0> sigma_raw;
}
~~~

~~~
transformed parameters {
   real EloStart[K];
   real<lower=0.0> k;
   for (i in 1:K) {
       EloStart[i] = EloStart_raw[i] - mean(EloStart_raw);
    }
   for (i in 1:K) {
       EloStart[i] = EloStart[i]/delta;
    }
   k = k_raw/delta;
   }
~~~

~~~
model {
   k_raw ~ normal(0, 1);
   sigma_raw ~ normal(0, 1);
   EloStart_raw ~ normal(0, sigma_raw);
   y ~ bernoulli(ProbFunction(EloStart, k, presence, N, K, Ai, Bi, delta));
   }
generated quantities{
   real<lower=0.0> sigma;
   sigma = sigma_raw/delta;
}
~~~

## S2 Update factor is equal to Logistic function

Using basic algebra, one can show that the update factor used in the definition of the Elo score is a “rather complicated written” form of the logistic function:

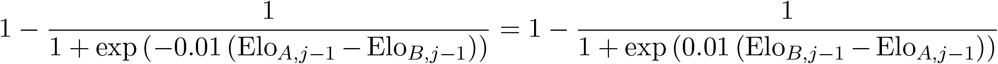

With *x* := 0.01 (Elo_*B,j*-1_ - E1o_*A,j*-1_), we further get:

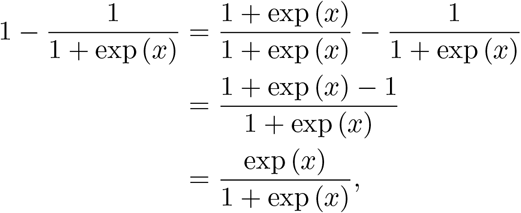

and therefore:

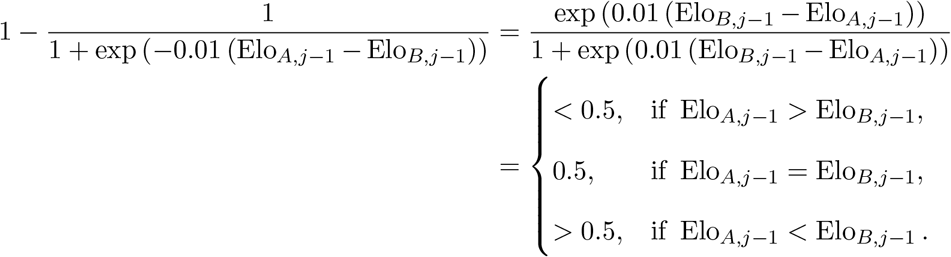

## S3 Standard deviation equivalence by varying Elo score difference factor *δ*

Let *X* = Elo_*A,j*-1_, and *Y* = Elo_*B,j*-1_, and further 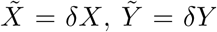, with 0 < ≠ = 1. In the case of equal variances 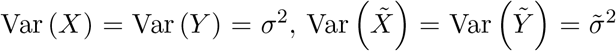, and independence, ie. 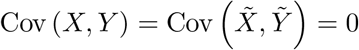:

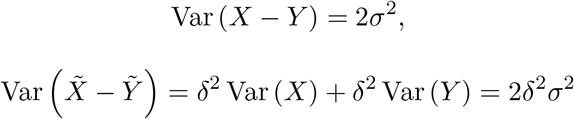

Since 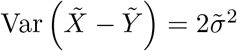, we get:

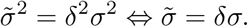

By re-defining *δ* = 1 in the winning/losing probability, differences in Elo scores that are 0. 01 as large as in the original definition with *δ* = 0.01 lead to the same winning/losing probabilities. Therefore, *k* will here also be only 0.01 as large as in the original definition, i. e. also the tax coefficient directly scales with *δ*:

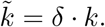

